# Opposing Strain Directions on Adjacent Left Ventricular Segments Predict Fibrotic Remodeling after Acute Myocardial Infarction

**DOI:** 10.1101/2023.03.20.533539

**Authors:** Tania Dubois-Mignon, Varouna Syvannarath, Marylou Para, Sylvain Richard, Pierre Sicard, Antonino Nicoletti, Giuseppina Caligiuri, Grégory Franck

## Abstract

**Background:** Despite similar levels of coronary occlusion and standard of care management, the occurrence of scarring over adaptive heart repair following acute myocardial infarction (AMI) remains unpredictable. Recent studies indicate that mechanical cues may modulate the transcriptional programs involved in tissue repair, possibly explaining why ventricular mechanical dyssynchrony an independent predictor of post-infarction outcome.

**Objective:** Our study aimed to investigate the relationship between mechanical cues and the outcome of post-myocardial infarction heart remodeling by live imaging. Specifically, we examined the impact of individual variability of myocardial dyssynchrony, characterized by a divergent direction of injured left ventricle wall movement next to live tissue, on the formation of a large scar, dilation of the left ventricle, and loss of pumping function.

**Methods:** We assessed the location and degree of regional systolic and diastolic dyssynchrony using transthoracic echocardiography coupled with speckle tracking imaging. Specifically, we measured the difference in absolute strain values between adjacent regions of the left ventricle at 5 days following the induction of a standard experimental infarction in female C57Bl6 mice. Three weeks later, transthoracic echocardiography was repeated to analyze the mass and global function of the left ventricle right before termination. We then examined the size of the scar in matched mid-sections of the left ventricle circumferential segments from each mouse using histomorphometry. Finally, we evaluated the potential impact on transcriptional tissue repair programs using spatial transcriptomic analysis on representative hearts with either adaptive or fibrotic post-infarction heart remodeling.

**Results:** We analyzed all 96 systolic and diastolic strain-related parameters in the same 48 regions of the left ventricle in all mice, with echocardiographic and histological sections following the same orientation. Stepwise analysis of the live imaging data revealed that a combination of 8 regional strain parameters could predict fibrotic remodeling (Area under the ROC curve= 0.8290). We observed that scarring remodeling was associated with opposing trends of systolic and diastolic circumferential strain % delta values on adjacent regions at day 5, while adaptive remodeling at day 28 occurred when the trend followed the direction of control (sham) hearts. Cluster analysis of gene transcripts and speckle tracking assessment on representative hearts with adaptive or fibrotic post-infarction remodeling indicated a correlation between regional post-infarction dyssynchrony and the transcriptional program. Adrenergic receptors, including *Adra1*, *Trpc3*, and *Myh7*, were found to be linked to specific regional dyssynchrony values and scarring remodeling.

**Conclusion:** Our study demonstrates the potential of regional strain parameters obtained through live imaging in predicting fibrotic remodeling following myocardial infarction. Furthermore, our findings suggest a link between regional post-infarction dyssynchrony and the transcriptional program. These results highlight the potential applicability of our approach in clinical settings and provide insights for future personalization of therapeutic strategies.

## Introduction

Ischemic heart disease is the most common heart disease and the leading cause of death in developed countries ^1^. It can manifest as an acute coronary syndrome which, when treated quickly by percutaneous revascularization, evolves favorably in the majority of cases. However, despite all the efforts for its management, post-infarction repair fails in up to 30% of patients who develop progressive post-ischemic heart failure ^2^. Understanding the pathophysiological mechanisms of unfavorable cardiac remodeling is crucial to developing new strategies to prevent or slow down disease progression.

The prognosis of survivors of acute myocardial infarction depends largely on the outcome of the subsequent cardiac remodeling ^3^. Hypertrophic remodeling of unaffected tissue is necessary to compensate for the loss of function in the ischemic area and maintain overall cardiac function, instead the generation of a large fibrotic sequela associated with left ventricle dilation can lead to terminal heart failure. Heart cells, such as contractile cardiomyocytes, vascular endothelial cells, smooth muscle cells, and interstitial fibroblasts and pericytes, must coordinate their actions and respond to biological and physical signals under physiological conditions. After an acute myocardial infarction, these cells must continue to fulfill their regular functions while compensating for the loss of function in a significant portion of the heart. Consequently, mechanical stresses within various regions of the heart become atypical, and the resident cells must adapt and respond to support contractility and repair the damaged tissue.

We hypothesized that specific atypical mechanical stresses could condition the outcome of myocardial remodeling and lead to fibrotic heart repair and progressive heart failure. To test this hypothesis, we aimed to capture both the mechanical cues and subsequent tissue responses at the same tissue localization. Decrypting the predictive value of such a putative mechanobiological coupling could provide valuable insights into the pathophysiology of cardiac remodeling and inform the development of new treatment strategies.

In recent years, there has been a significant advance in clinical imaging methods, allowing the acquisition of various motion parameters on a beating heart and facilitating the interpretation of mechanical forces. Previously, it was challenging to establish a map of local mechanical forces on a beating heart using non-invasive techniques. However, the development of recent clinical imaging methods such as magnetic resonance or ultrasound has made it possible to do so. One such method, speckle-tracking echocardiography, allows the precise evaluation of ventricle strain, strain rate, shear strain, and shear rate, which have been proposed as more sensitive tools to measure myocardial function. This advance is particularly important in understanding the pathophysiological mechanisms of unfavorable cardiac remodeling ^4, 5^.

From a molecular perspective, recent developments in spatial transcriptomics have allowed for an exhaustive and highly resolutive mapping of gene expression that helps define the mechanisms involved in a topological fashion ^6, 7^. By utilizing echocardiography in mice, we have developed a novel multidimensional imaging analysis that integrates speckle tracking measurements and transcriptomic profiling to assess the dynamic complexity of left ventricle (LV) post-infarction, allowing us to identify the mechanical stress measurement that is most likely to translate to the outcome of tissue repair in mouse post-infarcted hearts.

With the identification of the mechanical stress measurement that is most likely to correlate with tissue repair outcomes in experimental post-infarcted hearts, we aim to offer valuable insights that can guide the development of more efficient and personalized therapeutic interventions for patients suffering from ischemic heart disease.

## Method

### Mouse model of post-infarction remodeling

we placed a permanent ligation at the same level of the left coronary artery ^8^ in 20-week-old, female C57BL/6 mice, under anesthesia achieved by intraperitoneal injection of ketamine (100 mg/kg) and xylazine (5 mg/kg). The ligation creates prolonged ischemia, which is followed by a process of infarct expansion and resolution, fibrotic scarring, and the development of hypertrophy of the lateral left ventricle, allowing us to study “chronic” remodeling remotely from the infarction. We first After depilation of the thoracic area and local antisepsis with betadine, we intubated and ventilated the mice. To prevent hypothermia, we placed the animals on a thermostated pad (37°C) and monitored their body temperature using a rectal probe (**Fig. 1a**). We then performed a thoracotomy by delicately incising the 3rd left intercostal space. After separating the pectoral muscle and ribs, we incised the pericardium and sutured the left descending coronary under the microscope at the left atrioventricular junction. Finally, we closed and disinfected the mice and treated them with buprenorphine (0.05 mg/kg) and monitored them until they awakened.

**Figure 1:**
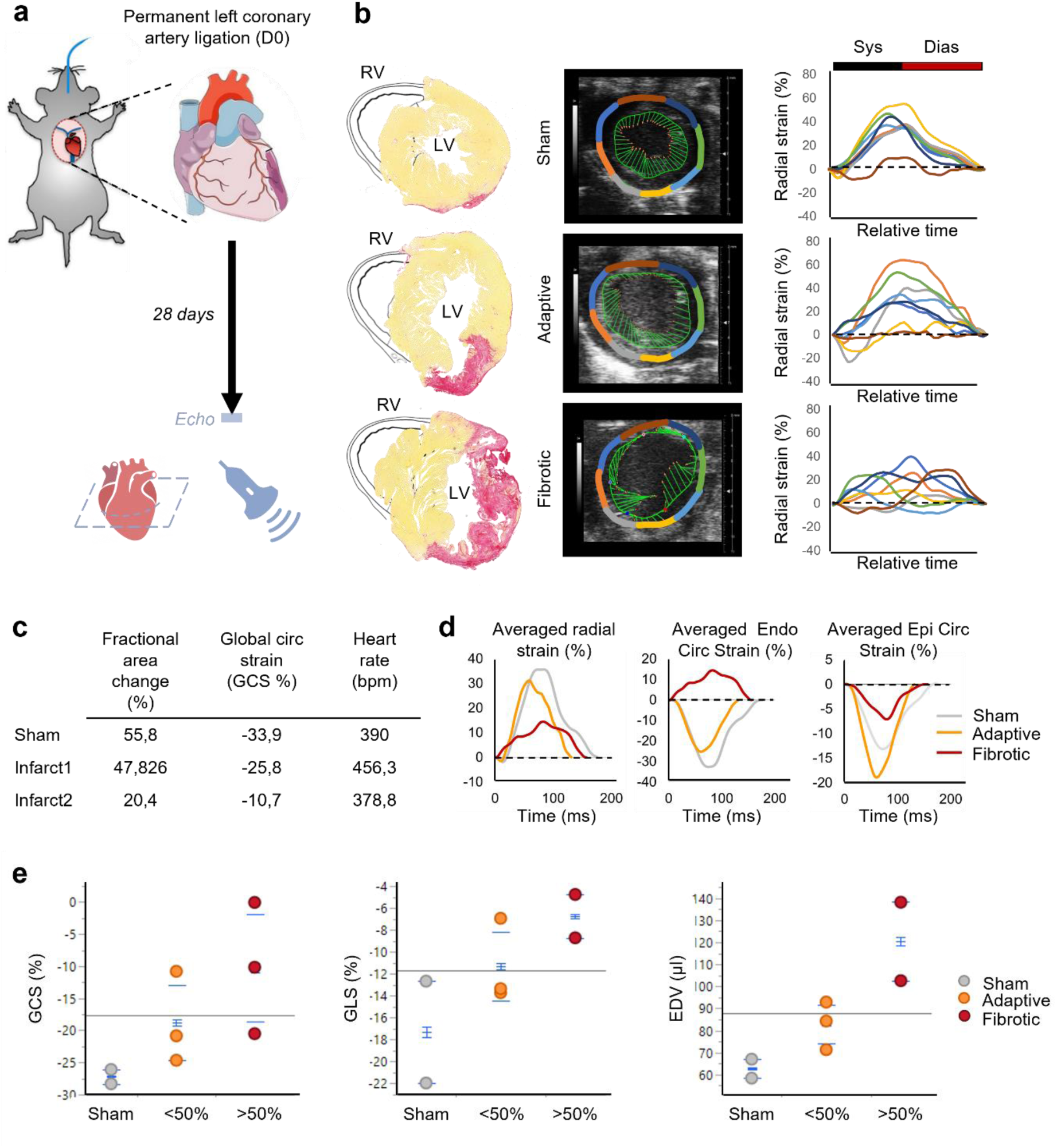
Global and segmental strain analysis used for group assignment of experimental mice. **a.** Experimental procedure from permanent left coronary artery ligation at day 0, to echocardiography at day 28. **b.** Radial strain map showing akinetic segments (yellow and brown, infarct 1; brown, grey, light blue, green, infarct 2), reflecting LV segments with deep contractile impairment. **c.** Values of fractional area change (%), global circumferential strain (%), and heart rate (bpm) shown for the 3 characteristics samples considered. **d.** LV-Averaged values of radial strain for each sample. **e.** Comparison of global longitudinal strain (GLS), global circumferential strain (GCS), and end-diastolic volume of the left ventricle (EDV) at day 28 in groups “sham”, “adaptive” or “fibrotic”.

### Transthoracic echocardiography and speckle tracking analysis

Five days after the procedure, we performed transthoracic echocardiography on anesthetized mice (isoflurane through a mouth-nose mask) placed on a heated pad (37°C). After heart rate stabilization, we used a 55 MHz ultrasound probe (MX550D MX, VisualSonics) and placed it on the depilated left hemithorax at a parasternal level and a short axis orientation. We acquired B-mode cine-loops with a total of 300 images during 3 entire heart cycles using a Vevo2100 echograph (VisualSonics). We traced the contours of the myocardial wall of the left ventricle (endocardium and epicardium) using the VevoStrain® software (VisualSonics). This method allows us to follow “natural” acoustic markers that are statistically evenly distributed within the tissue, and define 48 points of interest around the left ventricle whose displacement in space can be tracked over time (speckle tracking). For each of the 48 points, we analyzed six parameters, including velocity (point’s movement speed), displacement (point’s overall change in position), strain (percentage of deformation of the wall compared to its original shape), strain rate (speed at which strain occurs), shear (angular deformation between the epicardium and the endocardium), and shear rate (shear evolution over time). Positive values of strain are usually regarded as elongation, while negative values translate myocardial shortening. In this study, we considered an increase or a decrease in the absolute value of strain to evaluate net elongation. Since the contraction of the left ventricle is a complex phenomenon involving thickening, shortening, and torsion, we acquired some of these parameters in two perpendicular axes: circumferential (shortening/elongation) and radial (thinning/thickening of the wall, **Fig. 2a**). We also evaluated the temporal values for each parameter by taking into account mean values, peak values (amplitude), and the time at which the peak value was reached (time to peak, TTP). Accordingly, 96 parameters were obtained for each of the 48 LV points, representing 4608 dynamic data point per mouse heart. Statistical analysis was performed using JMP® software (SAS Institute Inc). Nominal logistic regression was applied to all raw data, and stepwise regression was used to identify combinations of parameters to build the receiver operating characteristic (ROC) curve (with positive response indicating fibrosis). Two-way ANOVA was conducted to assess differences between adaptive and fibrotic remodeling in terms of delta strain on adjacent left ventricle regions.

**Figure 2:**
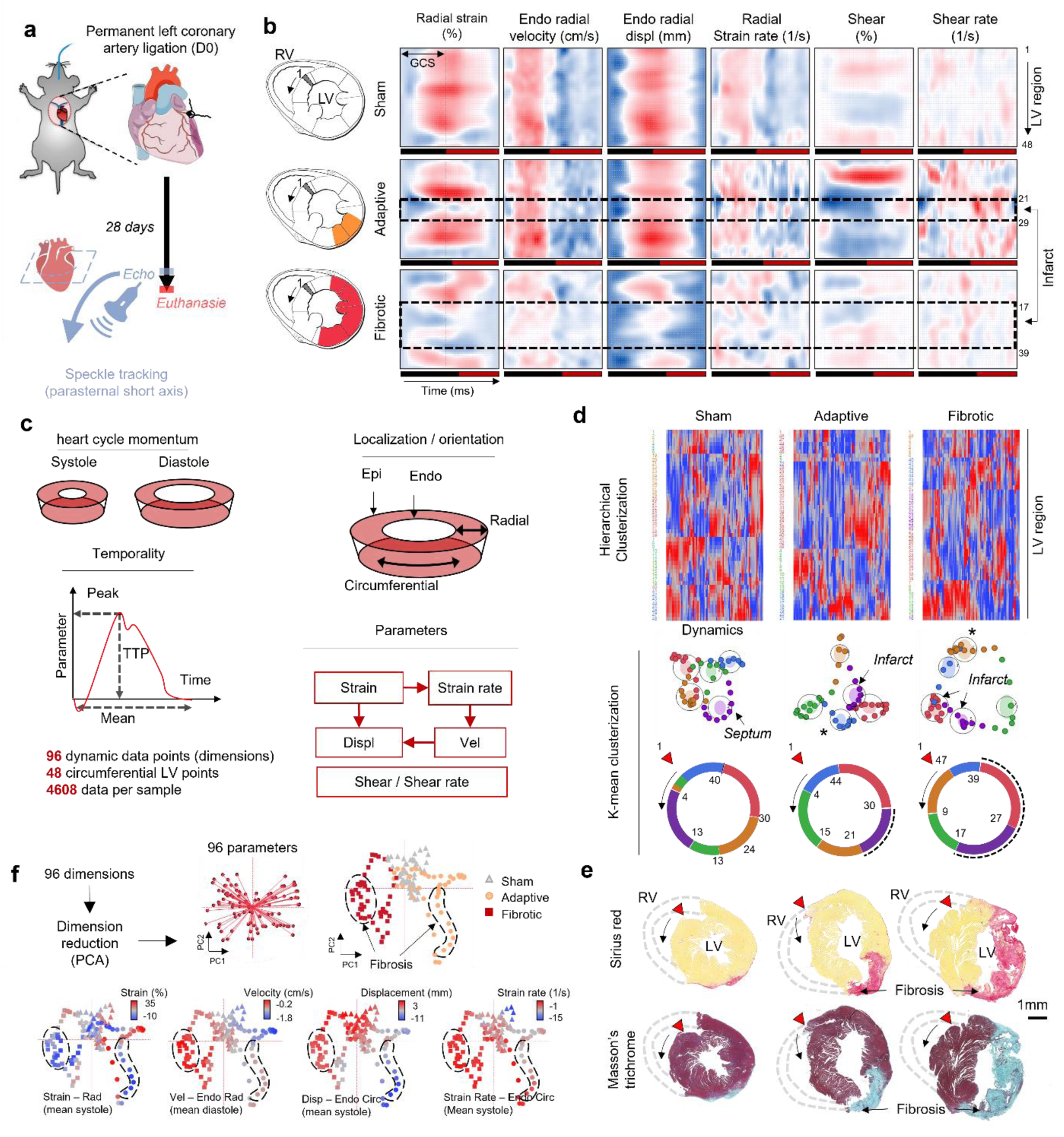
Comprehensive evaluation of ventricular regional mechanic cues in sham and post-infarction hearts using multiparametric speckle-tracking analysis. **a.** Experimental procedure. **b.** representative examples of heart motion heat maps showing heterogenous motion profiles in sham, adaptive, and fibrotic hearts, 28 days post infarct. **c.** Motion parameter clipping according to heart momentum, localization, orientation, and temporality, for strain, strain rate, displacement, velocity, shear, and shear rate. **d.** Heatmaps and dimension reduction by principal component analysis (PCA) showing intra-ventricular motion heterogeneities, superimposing with histology performed on transversal heart sections of the corresponding areas (**e**). **f.** Inter-ventricular heterogeneities showed by PCA and K-means clusterization, and plotted against various motion parameters determined in **c**.

### Histology and correlation between biological and mechanical parameter

After the echocardiography, mice were terminated by isoflurane overdose, and the hearts were excised, fixed in 4% paraformaldehyde at 4°C for 48 hours, and embedded in paraffin. Six µm-thick cross-sections were obtained from the corresponding parasternal short axis. Areas of fibrosis were identified by Sirius red and Masson’s trichrome staining, and regions exhibiting kinetic disorders in the echocardiography loops (hypokinetic, akinetic, or dyskinetic) were used to manually overlay the parameters obtained from the ultrasound images and histology (**Fig. 1c, 2b and 2e**).

### Spatial transcriptomics

Three histological FFPE sections from mouse hearts were used for spatial transcriptomics analysis (HistIM platform, Cochin Institute, https://support.10xgenomics.com/spatial-gene-expression). Hearts with no infarct (sham), small, or large infarcts were selected. The FFPE sections were mounted on Visium slide capture areas (6.5 x 6.5 mm, 10X Genomics), which contained 5,000 spots of 55 µm diameter, spaced 100 µm apart (**Supplementary Fig. 1c**). Each spot was composed of oligonucleotides with a unique spatial barcode. Hematoxylin and eosin staining were performed to retrieve precise heart morphology after deparaffinization. The antigen de-crosslinking step was performed by warming the slides at 70 °C for 1 hour, releasing sequestered mRNAs. A cocktail of primers that specifically hybridized to virtually all the murine mRNAs was then added to the tissue. An RNAse treatment (PN-3000605, Visium FFPE Reagent Kit, 10X Genomics) and a permeabilization step (Perm Enzyme B PN-3000602, Visium FFPE Reagent Kit) were performed to foster primer hybridization to the spot-specific oligonucleotide barcodes. Hybridized primers were carefully collected from the slides and used for quantitative PCR (qPCR) to determine the number of PCR cycles for each sample. Indexed primers were added to group several samples (Dual Index Plate TS Set A, 10X Genomics). Then, DNA fragments were purified using SPRIselect (Beckman Coulter) to prepare the cDNA library. Finally, Illumina paired-end sequencing was performed (Genomic’IC platform, Institut Cochin), and the sequencing depth averaged 62,500 reads per spot.

### Bioinformatics analysis

The spatial transcriptomic data was visualized using Partek® Flow® genomic analysis software. The Space Ranger 1.3.0 algorithm was first applied to align sequences and create a matrix from the spatial barcodes. The resulting data was then superimposed on histology (**Fig. 5**). Spots with insufficient information were filtered out, as well as genes detected in less than 1% of the spots (**Supplementary 1d**). A SC Transform normalization (Seurat) was applied to account for variable sequencing depth across spots ^9^. The Seurat3 integration function was then used to normalize inter-sample variability. Principal component analysis (PCA) was performed to classify spots by expression profile, followed by Uniform Manifold Approximation and Projection (UMAP) and graph-based unsupervised clusterization. Gene expression analysis between clusters was carried out using gene-specific analysis (GSA), using the best statistical model to calculate p-values and regulation factor (fold change). Gene Ontology (GO) term enrichment of the differentially expressed genes (DEGs) was performed using statistical overrepresentation test (Fisher’s exact test with Benjamini-Hochberg false-discovery rate [FDR] multiple-test correction) using GESEA. The spots were then pooled into 24 circumferential LV segments, and the mean mRNA expression for each segment was compared to the superimposed mechanical data obtained by echocardiography (**Fig. 6a**).

## Results

### Regional strain analysis by speckle-tracking in post-infarction hearts as a tool for predicting unfavorable post-infarction heart remodeling

The functional outcomes of the experimental hearts on day 28 varied considerably despite similar coronary ligation levels (**Fig. 1a-c**). To determine the attribution of mice to different groups, we first divided the experimental mice based on the extent of fibrotic sequela occupying more than 50% or less than 50% of the left ventricle (n=3 in each group). Based on the extent of fibrosis in heart sections histomorphometry at day 28, mice were therefore assigned to the “adaptive” (<50%) or “fibrotic” (>50%) remodeling groups. The analysis of strain parameters over a complete heart cycle (relative time covering both systole and diastole) revealed reduced radial strain and opposing circumferential strain percentages in the fibrotic remodeling group compared to sham and adaptive remodeling groups, reflecting the functional failure of post-infarction remodeling (**Fig. 1b-c**). Analysis of global function, as assessed by global longitudinal and circumferential strain (GLS and GCS), clearly demonstrated that functional long-term recovery was associated with a reduced extent of the fibrotic sequela (**Fig. 1d**). Additionally, the end-diastolic volume (EDV) of the left ventricle was consistently higher in mice of the fibrotic remodeling group than in mice with adaptive remodeling, reflecting the dilation of the failing heart and further supporting the classification used for comparing the two types of remodeling outcome. (**Fig. 1e).**

### Multiparametric heart motion analysis by speckle-tracking echocardiography

Evaluation of velocity, displacement, strain rate, shear, and shear rate is possible, although their interpretation separately is challenging, if not impossible. Each parameter was taken independently and harbored time and regional variability (**Fig. 2b** and **Supplementary Fig. 2-3)**. In addition, we analyzed an exhaustive set of 96 dynamic parameters derived from speckle-tracking exploration, including heart cycle momentum (systole, diastole), localization (endocardial or epicardial), orientation (circumferential, radial), and temporality (mean value, peak amplitude, time to peak) for each of the six studied parameters (velocity, displacement, strain, strain rate, shear, shear rate, **Fig. 2c**).

Taking into account these 96 parameters, we could observe important heterogeneity along the 48 LV segments using hierarchical clusterization, and a dimension reduction by PCA followed by a K-means clusterization (**Fig. 2d**). Notably, this clusterization closely superimposed with histological features highlighted by Sirius Red and Masson’s trichrome staining, such as infarcted areas (**Fig. 2e**). Finally, we could identify important inter-sample specificities using a PCA on the multiparametric motion data, followed by K-means clusterization, when comparing sham, adaptive and fibrotic hearts (**Fig. 2f**). In both types of infarcted heart, we observed a divergence starting from a physiological contractile phenotype superimposing with sham (grey points), to LV regions with fibrosis (dashed lines). Notably, while strain values well delineated intra-sample infarct areas with low strain found in infarcted area, each infarct harbored opposite specificities regarding velocity, displacement, and strain rate, identifying inter-sample variabilities (**Fig. 2f**).

### Multiparametric heart motion echography as a tool for predicting unfavorable post-infarction heart remodeling

After exploring different model types, the statistical analysis software determined that a nominal logistic fit was the most appropriate. The analysis identified 8 parameters that were potentially associated with fibrosis (**Fig. 3b**). To further evaluate the predictive value of these parameters, we conducted a stepwise regression analysis and found that the combination of all of them had the greatest predictive power for fibrosis. The resulting ROC curve indicated that the combination of these parameters was able to predict the occurrence of fibrosis at day 28 with an area of 0.8290 (**Fig. 3c**). This suggests that the combination of these 8 parameters is a strong predictor of the development of fibrosis over time. These observations highlight deep dynamic specificities and point to a strong influence of post-infarction mechanical cues on the fate of heart repair.

**Figure 3:**
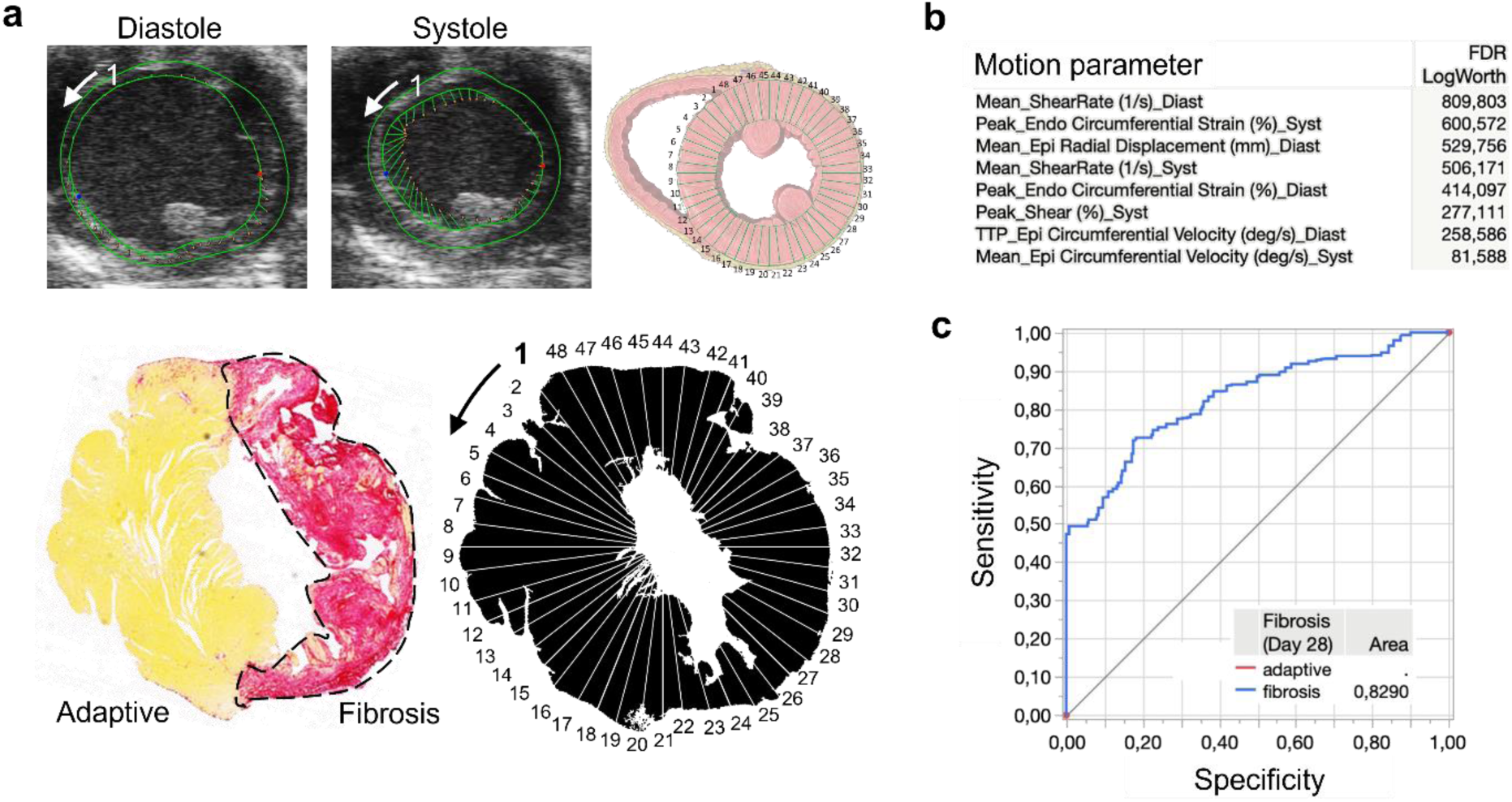
Selected motion parameters at day 28 predict unfavorable post-infarction LV remodeling. **a.** Regional segmentation shown in a representative fibrotic heart post-infarction. **b.** Motion parameters significantly associated with fibrosis occurrence by nominal logistic fit, and resulting receiver operating characteristic curve (**c**).

The main objective of our study was to assess the atypical cues generated by the divergent direction of adjacent segments in post-infarction hearts that could serve to identify the unfavorable mechanobiological coupling at the regional level. To achieve this objective, we calculated the delta between the 48 values for each of the identified 8 parameters in all experimental mice (n=6) as well as in 2 control shams.

After comparing the values of each of the 8 strain parameters across the 48 adjacent segments of the left ventricle, we observed a trend in experimental mice with adaptive post-infarction remodeling that closely paralleled that of the sham controls for the trend line of circumferential strain % deltas. Conversely, in experimental mice exhibiting fibrotic-dilative remodeling, we observed an opposite trend, represented by a sine wave, in both systole and diastole (**Fig. 4**)

**Figure 4:**
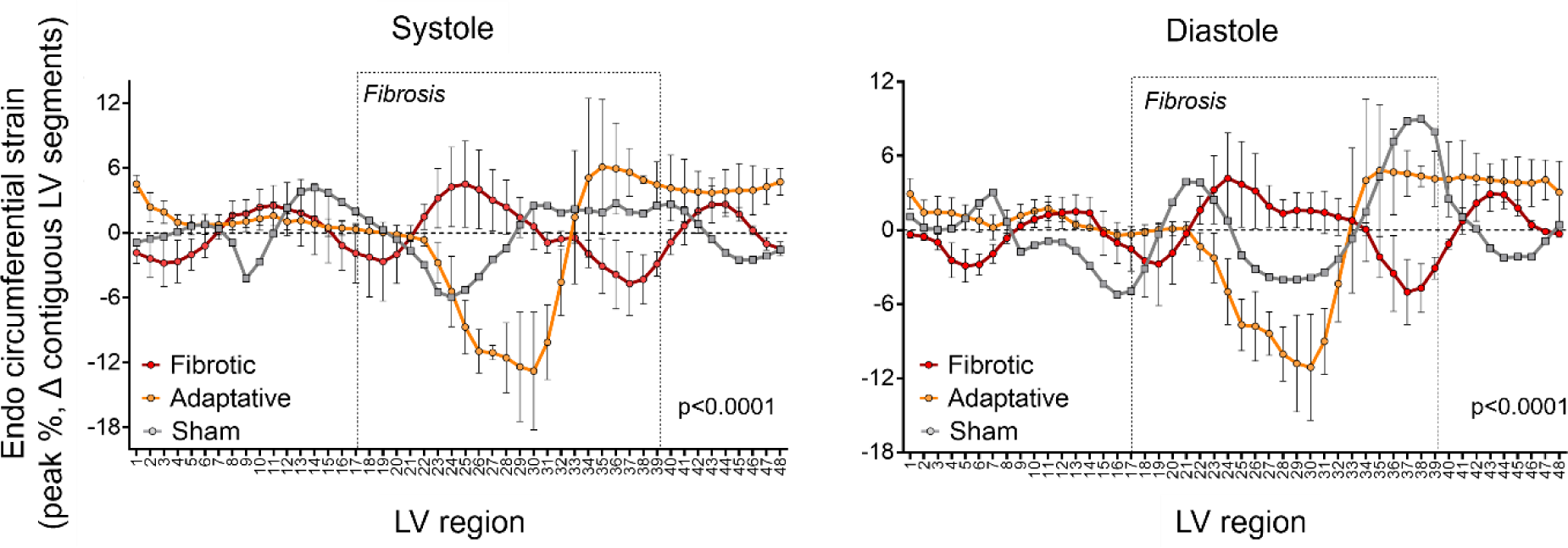
Left ventricle motion dyssynchrony at day 5 assessed by delta peak endocardial circumferential strain. Systole and diastole motion are shown for shams (grey), adaptive remodeling heart (orange), and fibrotic remodeling hearts (red). Differences were analyzed by two-factor ANOVA (LV region x group; p<0.0001).

### Exploratory spatial transcriptomics suggest distinct transcriptional signatures in adaptive and fibrotic remodeling post-infarction hearts

To investigate the putative biological relevance of the opposite mechanical strain on transcriptional programs during post-infarction heart repair, we combined strain analysis with spatial transcriptomic analysis (**Fig. 5a**). Visium was used to orient, process, and analyze corresponding heart sections, and principal component analysis followed by graph-based clustering revealed areas of specific signature (**Fig. 5b**). Differential expression of each cluster compared to sham was shown using unsupervised hierarchical clustering and revealed nodes of highly significantly regulated genes (**Fig. 5c and d**).

**Figure 5:**
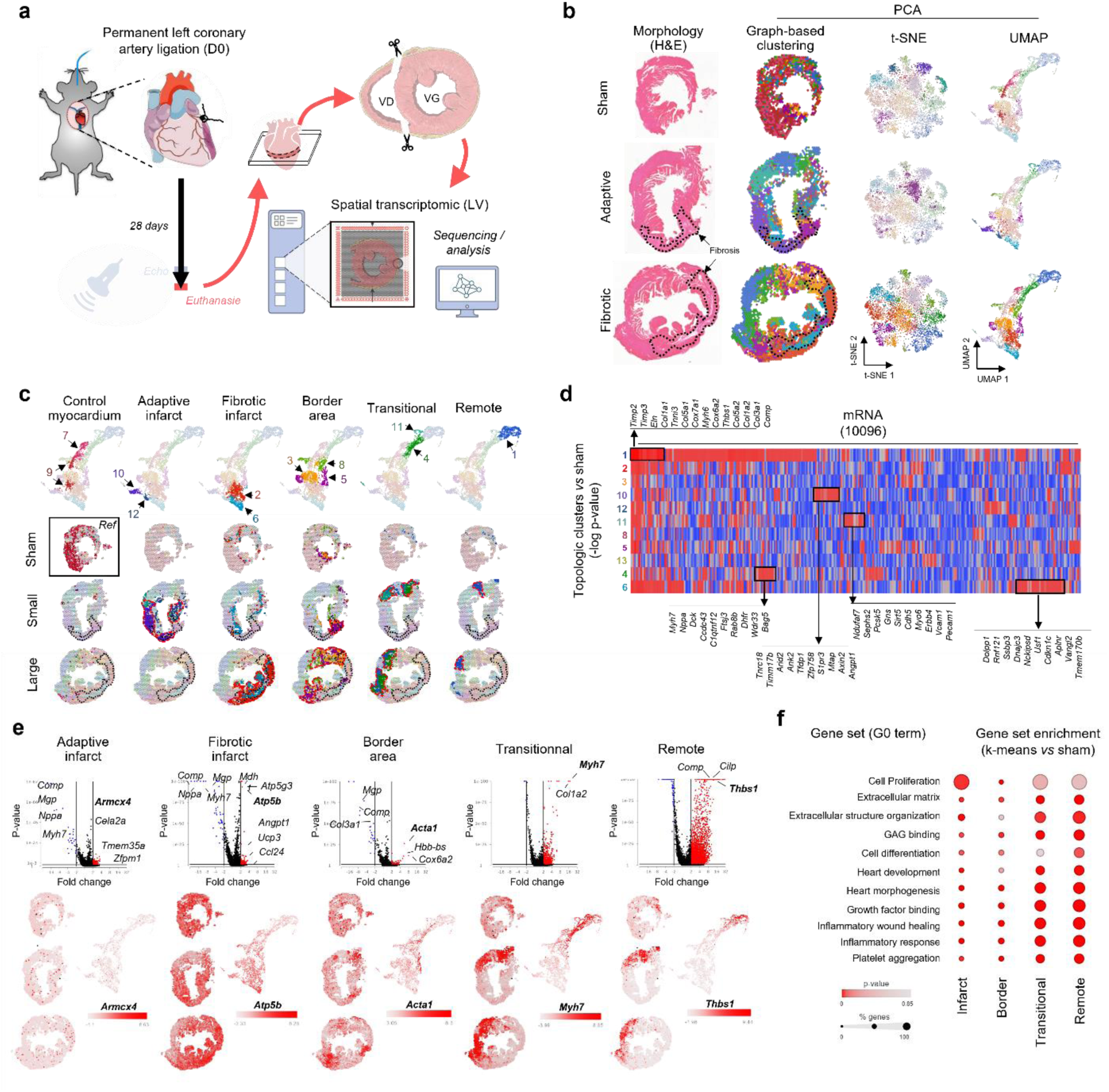
Spatial transcriptomics analysis in adaptive and fibrotic remodeling post-infarction hearts. **a.** Visium (10X genomics) was used to orient, process, and analyze corresponding heart sections. **b.** Principal component analysis followed by graph-based clustering revealed areas of specific signature, which were superimposed on heart histology. Both t-distributed stochastic neighbor embedding (t-SNE) and uniform manifold approximation and projection (UMAP) plots are shown with clusters. **c.** Both adaptive and fibrotic infarcts harbored a transcriptomic gradient shown using UMAP, going from the infarct area to the remote zone, passing through border zones. **d.** Differential expression (-log p-value) of each of the 13 clusters compared to sham was shown using unsupervised hierarchical clustering and revealed nodes of highly significantly regulated genes (p<0.01). **e.** Clusters were pooled to form specific key regions according to histology, and whole mRNA expression within each group was compared to sham heart section mRNA expression. Differential analysis for each comparison showed hundreds of regulated genes, with the highest regulation found in both transitional and remote areas. *Acta1* was overexpressed in the border area, and *Myh7* was highly overexpressed in both transitional and remote areas. **f.** For each cluster, DEGs compared to sham drove a GO term enrichment. GO terms related to heart development, heart failure, and extracellular matrix were found enriched in DEGs within “transitional” and “remote” areas.

Sham heart section showed a relatively homogeneous signature, whereas both small and large infarcts harbored a transcriptomic gradient going from the infarct area to the remote zone, passing through border zones. Clusters were pooled to form specific key regions according to histology and referenced as “adaptive infarct specific,” “Fibrotic remodeling specific,” “Border area,” “Transitional,” and “Remote.” Whole mRNA expression within each group was compared to sham heart section mRNA expression (**Fig. 5e**). Differential analysis for each comparison showed hundreds of regulated genes, with the highest regulation found in both transitional and remote areas (**Fig. 5f, Supplementary table**).

### Correlating Mechanical Forces and Myocardial Transcriptomics in Post-Infarction Hearts

To evaluate the relationship between mechanical forces and time-matched myocardial transcriptomic one month post-infarction, we superimposed ultrasound data from day 28 with the spatial transcriptomic analyses, in 24 circumferential regions of interest for each sample **(Fig. 6a**). The analysis revealed a significant coupling between mRNA expression and mechanical parameters for 3538 mRNA (32%) in the large infarct, 1542 mRNA (14%) in the small infarct, and only 560 mRNA (5,1%) in the sham heart (**Fig. 6b**). In the large infarct, 28/96 parameters (30%) were significantly correlated with more than 5% of the mRNA set (**Fig. 6d**). Displacement, velocity, strain and strain rate constituted the significant metrics in sham and small infarcts, with similar respective ratios. In the large infarct, significant coupling with strain rate was increased, and shear / shear rate was also found meaningful (**Fig. 6c**).

**Figure 6:**
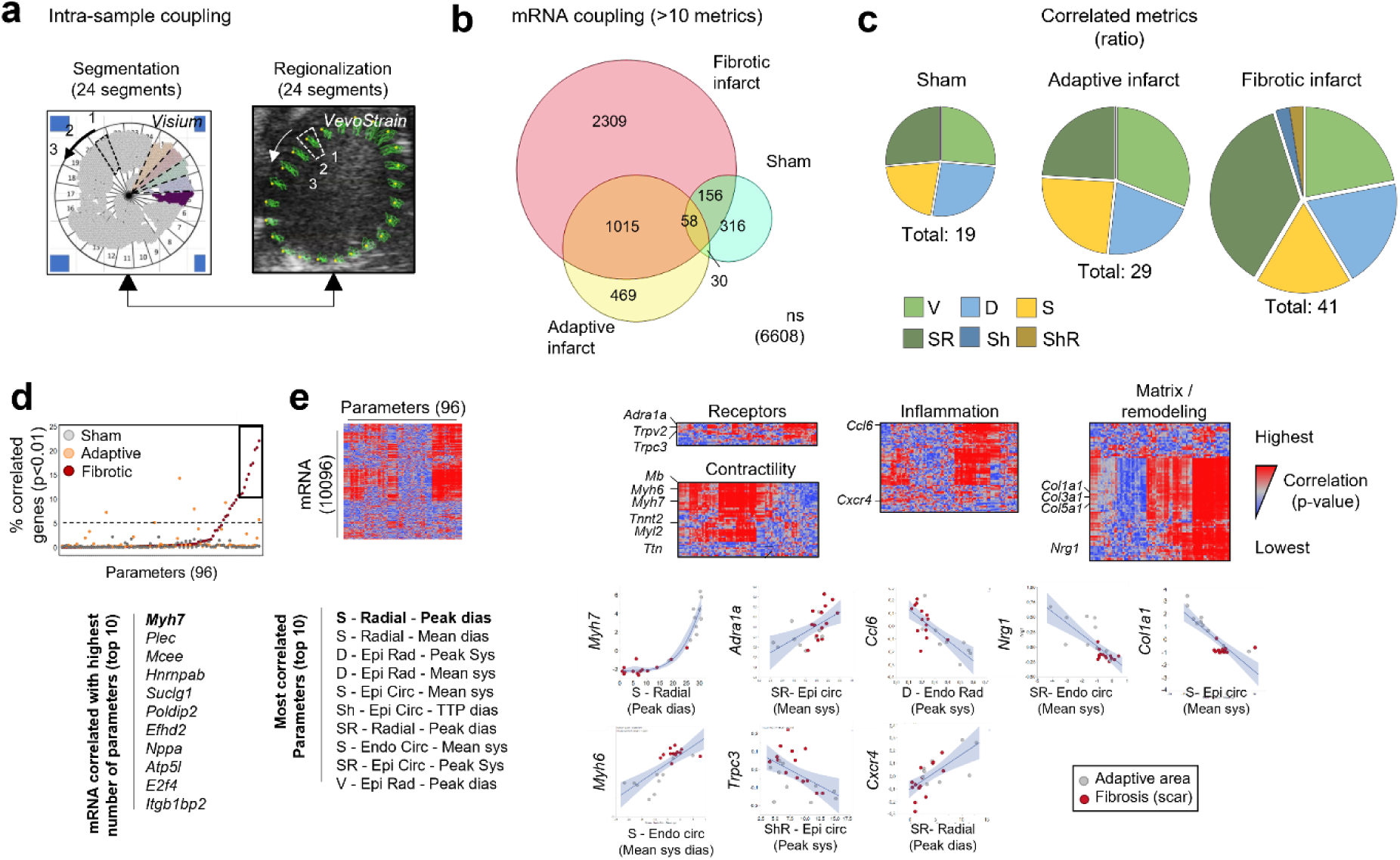
Correlation between myocardial transcriptomic analysis and mechanical forces on post-infarction remodeled hearts. **a.** Spatial organization of the circumferential regions of interest generated from superimposing ultrasound and spatial transcriptomic analyses. **b.** Percentage of correlated mRNA with mechanical parameters for each sample type. **c.** Significant mechanical parameters in each sample type. **d.** Percentage of parameters significantly correlated with mRNA in a fibrotic infarct. **e.** unsupervised hierarchical clustering of the whole mRNA set according to the 96 parameters.

Among the mRNA correlating the most with mechanical parameters, *Myh7*, *Nppa* and *Atp5l* were overexpressed in transitional and remote areas as compared to sham. To classify the whole mRNA set according to the 96 parameters, unsupervised hierarchical clustering revealed nodes of thousands of genes specifically correlated with a subset of parameters (**Fig. 6e**).

We delimited this study to gene sets relevant to pathological myocardial remodeling, including heart receptors, inflammation, and matrix / remodeling. The relationship between mRNA expression and parameters in the gene set “matrix / remodeling” appeared more complex, as non-linear regression was generally observed, with low mRNA expression values observed in infarcted segments. Coupling data from the adaptive infarct revealed a different profile, with specific genes and parameters illustrated in **Supplementary Fig. 9.** Radial strain (diastolic peak) was the only common parameter found among the top 10 associated parameters in both infarct types. This indicates that the number of correlated mRNA decreased with infarct size (**Fig. 6b and 6d**).

### Linking Multiparametric Strain Metrics to Gene Expression Profiles Reveals Dynamic Polarization of LV Segments

To investigate the link between the dynamic LV phenotype shown by PCA and the biology of such LV segments, we classified and pooled the 24 circumferential segments for each sample based on both a PCA classification using the echo parameters (**Fig. 7a and 7b**) and a projection (UMAP) using the transcriptomic data (**Fig. 7c, left**) or a pool of genes of interest (GOI, n=250) used to highlight contractility, matrix-remodeling, and receptors gene sets (**Fig. 7c, right**). K-means clusterization of both UMAP identified circumferential segments with distinct signatures. Surprisingly, this characterization strongly superimposed, whether the transcriptome data or the GOI were used. Overall, this analysis shows that the expression of a small panel of genes critically involved in myocardial functions is intimately linked to multiparametric echo metrics.

**Figure 7:**
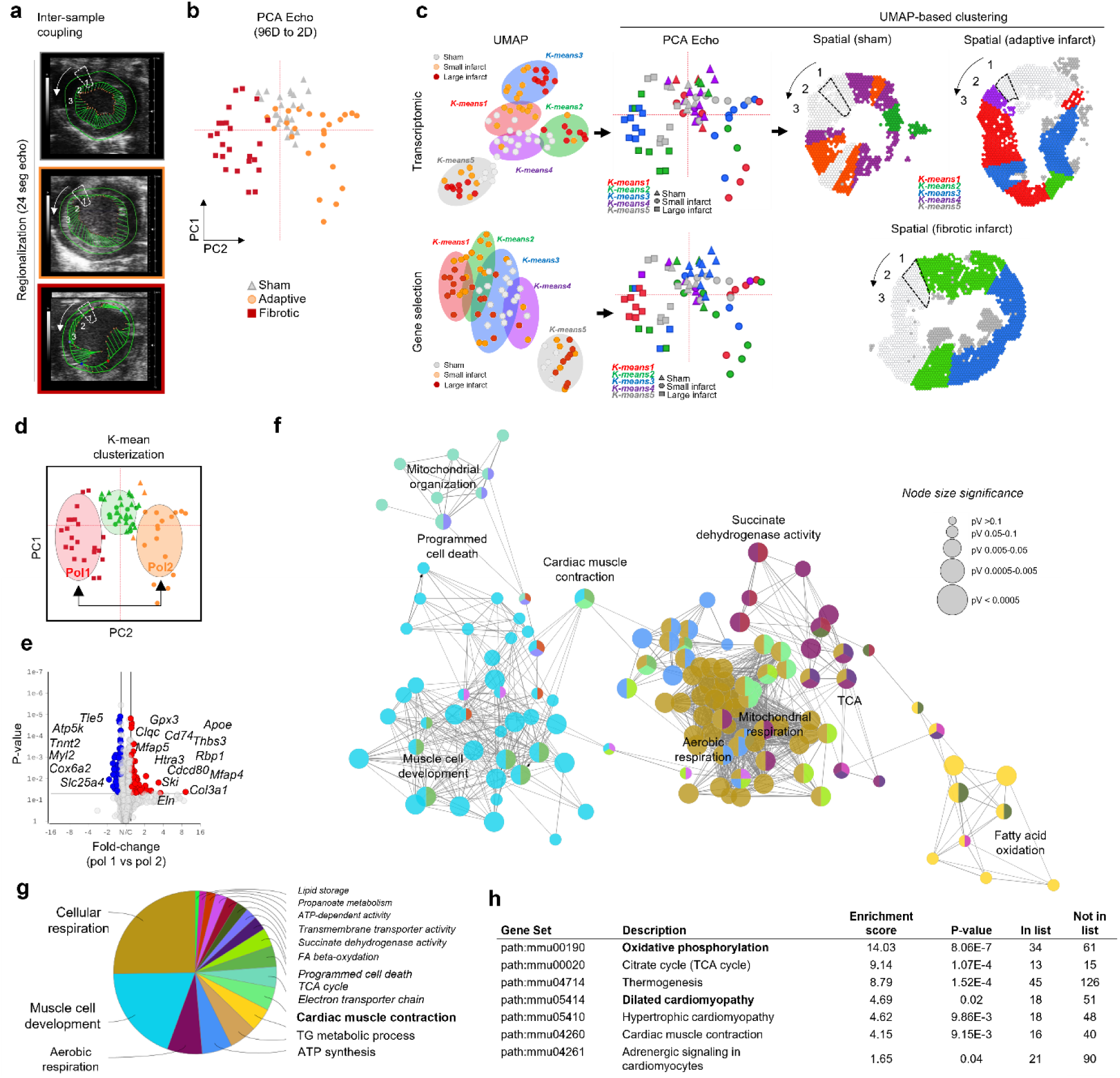
Classification and clustering of LV segments based on both echo parameters and transcriptomic data. **a-b** Principal component analysis (PCA) classification of LV segments using echo parameters. **c.** Uniform manifold approximation and projection (UMAP) classification using transcriptomic data (top) or a pool of genes of interest (GOI, bottom). K-means clusterization of both UMAP identified circumferential segments with distinct signatures. **d.** Dimension reduction of motion parameters by PCA, and K-means clusterization of LV segments, where two groups (“Pol1” and “Pol2”) distant from LV points of physiological contractility (green points) and superimposing to sham points were compared. **e.** Two-way ANOVA test comparing mRNA expression profiles between “Pol1” vs “Pol2” and the resulting up-regulated and down-regulated mRNA, with fold change >1.5. **f-g** Pathway enrichment analysis using ClueGO integrating KEGG/Biocarta pathways. **h.** Top 25 KEGG pathways retrieved from the analysis, including metabolic pathways (oxidative phosphorylation, TCA cycle, and thermogenesis), cardiomyopathies, and cardiac physiology (dilated cardiomyopathy, hypertrophic cardiomyopathy, cardiac muscle contraction, and adrenergic signaling in cardiomyocytes).

To identify groups of LV segments harboring dynamic polarization, we performed K-means clusterization (**Fig. 7d**). Two groups (“Pol1” and “Pol2”) distant from LV points of physiological contractility (green points) and superimposing to sham points were compared. A two-way ANOVA test comparing mRNA expression profiles between “Pol1” vs “Pol2” was performed, and revealed 156 up-regulated (e.g. *Tle5*, *Atp5k*, *Tnnt2*, *Myl2*, and *Cox6a2*) and 143 down-regulated mRNA (e.g. *Col3a1*, *Eln*, *Mfap4*, *Apoe*, and *Thbs3*, **Fig. 7e**).

These results demonstrate that an imaging-based analysis of heart motion provides important information regarding ventricle function beyond a mechanical readout.

## Discussion

Our study investigated the use of non-invasive medical imaging, specifically regional strain analysis using multiparametric speckle-tracking, to identify individuals at risk of unfavorable post-infarction heart repair as early as 5 days after a myocardial infarction. We found that this method directly correlates mechanical stress with biological function and offers a more accurate and sensitive classification for clinical cardiology. While a multi-parametric approach has not yet been widely adopted due to technological limitations and the computationally demanding nature of analyzing complex datasets, we anticipate that as technology and software continue to improve, more studies exploring multi-parametric approaches will emerge in the near future, ultimately improving the diagnosis and prognosis of cardiopathies.

Our exploratory spatial gene expression analysis revealed a strong association between intra-ventricular dynamics and transcriptomic signature involving mRNA encoding contractile and synthetic proteins. Surprisingly, we found that the mRNA expression of a broad variety of collagens was strikingly and exclusively upregulated in the remote zones within the non-fibrotic tissue opposed to the infarction, indicating active remodeling, which was undetectable using conventional histology. Our study provides insights into the remodeling process and highlights the need for a more comprehensive understanding of the cellular and molecular mechanisms underlying extracellular matrix (ECM) remodeling following myocardial infarction. Traditional histological analysis alone may not be sufficient to capture the complexity of the ECM remodeling process, and future analysis of ultrastructural modifications could help distinguish the precise cardiomyocyte phenotype and matrix remodeling state related to stress.

We also found evidence to suggest a role for mechanobiology on mRNA encoding receptors engaged by heart deformation. We observed a significant effect of heart deformation on the mRNA encoding receptors targeted by the current treatments against heart failure, including α- and β-adrenergic receptors, which are known to be affected by strain ^10^. We also found that α- and β-adrenergic receptor expression carries a cell specificity that is likely affected by stress indirectly ^11^. Additionally, the expression of numerous mechanoreceptors, such as the transient receptor potential (TRP) family transcripts, was associated with myocardial deformation. The complexity of the signaling pathways involving these receptors and mechanoreceptors highlights the need for further investigation to understand their precise role in heart deformation and the potential for new therapeutic targets. Our findings shed light on the intricate molecular mechanisms underlying heart function and suggest new avenues for developing targeted therapies for heart failure. Although the expression of TRPs and adrenergic receptors in cardiomyocytes and cardiac fibroblasts has been extensively studied ^12–16^, their occurrence in other cardiac cells such as endothelial cells (ECs) or leukocytes is still limited. Given that ECs constitute around 10% of the cell population in the physiological myocardium and express a wide variety of receptors, including TRPs, it is possible that mechanoperception of atypical stress by ECs could be a key pathophysiological parameter linking stress, remodeling, and heart failure.

Our study investigated the expression of neuregulin-1 (NRG-1), an embryonic epidermal growth factor almost exclusively expressed by ECs in the myocardium. We found that *Nrg-1* is strongly upregulated in the myocardium opposed to the largest infarcts and harbored an expression pattern significantly correlated with 22 motion metrics, including radial strain, radial and circumferential strain rate, radial epicardial velocity, and radial endocardial displacement. In vitro, *Nrg-1* expression was increased in microvascular ECs subjected to cyclic stretch (**Supplementary Fig. 8**). However, it was not regulated in the remote area facing the small infarct, suggesting that mechanical stress must reach a certain threshold to switch on *Nrg-1* expression, which was not reached in this region.

Our study also highlighted a high degree of inter-individual variability, and it is noteworthy that variabilities correlated with transcriptional specificities, indicating that discreet myocardial biology may be interpreted by noninvasive imaging alone. We provide a proof of concept that early multidimensional stretch imaging can predict the outcome of heart failure, as assessed by ejection fraction, fractional area change, and fibrosis extent. Using this approach, an exhaustive inter-sample characterization using dynamic imaging could help define patient profiles with important prognostic value.

The expression regulation of mRNA encoding for clinically targeted receptors, as well as the modification of their downstream signaling pathways, suggest that dynamic imaging could be a valuable tool in predicting patient response to treatments. However, larger experimental cohorts will be necessary to draw more definitive conclusions. A multi-centered database of patient information could be investigated to establish classification and learning models that could aid in predicting outcomes. Additionally, human heart biopsies obtained after patient transplantation may provide valuable insight in bringing this concept to the clinic.

### Study limitations

Although speckle tracking is a more sensitive method than LVEF to measure heart contractility, it is important to note that strain is influenced by changes in cardiac load and hypertrophy ^17^. Strain rate, which is less influenced by these confounders, has been found to be robust and able to predict discrete and early functional alterations in mouse models of MI ^18^. The use of radial shear has also provided interesting conclusions ^5^. However, all of these approaches carry technical limitations inherent to the technique of 2D speckle tracking, including stationary image and off-plane artifacts, or the absence of “out-of-plane” speckle motion that may compromise reliability in a clinical context ^19, 20^. Sources of variation also exist in speckle-tracking based analysis, including the echocardiographic views obtained for image acquisition, which can result in capturing different fiber layers at different levels and, thus, yield variable results. This is especially important for short-axis in our case, since we aim at superimposing echo and spatial transcriptomic data. However, single plan measurements are still helpful in the post-MI model, as clear anatomical changes resulted from remodeling and drove serial delineation. Investigating 4D strain loops may greatly help to overcome this problem. Finally, LV tracing performed by the investigator can constitute another source of variability as stretch values depend on the quality of the tracing. This variability should be minimized through the use of a standardized laboratory protocol.

In addition, the spatial transcriptomic approaches used in this study have limitations in assessing the cell contribution to the transcriptomic signature because the resolution is limited to the spot size (55 µm diameter, 100 µm apart), generally enriching 1 to 10 cells. Further studies should either use parallel single-cell evaluation or embrace the use of spatial transcriptomic technologies at cell and subcellular resolution. Paired single-cell and spatial transcriptomics could also help map the cell interactome. While these technologies are already available, their cost is still very high today. It is also important to note that transcriptomic evaluation, while translating regulation of exhaustive signaling pathways, upstream regulators, or transcription factors, never assesses protein expression, which constitutes the most relevant evaluation of biological changes. By using conventional protein immunostaining or unbiased spatial spectroscopy, identification of local tissue protein regulation should constitute an important exploratory work in the future.

Furthermore, the transcriptomic data in this study were only obtained in a subset of experimental mice, and the sample size was limited. Therefore, the results of this section should be considered exploratory. Although the data do suggest a strong link between intra-ventricular dynamics and transcriptomic signature, further studies will be needed to confirm and expand upon these findings. It is also essential to replicate these results in larger cohorts and in other animal models or human patients to validate the significance of the link between LV dynamics and gene expression. Overall, our study provides an initial proof of concept and highlights the potential value of transcriptomic data in understanding the underlying mechanisms of heart disease.

### Conclusion

This study offers a groundbreaking level of detail in uncovering biological responses in the myocardium. While it has established important correlations between biology and motion metrics, further research is necessary to establish causality. Exciting future experiments, such as in vitro cell culture under varying stretch conditions or in vivo studies utilizing knockout mice or specific inhibitors, may offer crucial insights and validate the targets identified in this study. With its important findings, this study highlights the potential for dynamic imaging to guide treatment decisions and predict outcomes in cardiac patients.

## Supplementary Figures

**Supplementary Figure 1:**
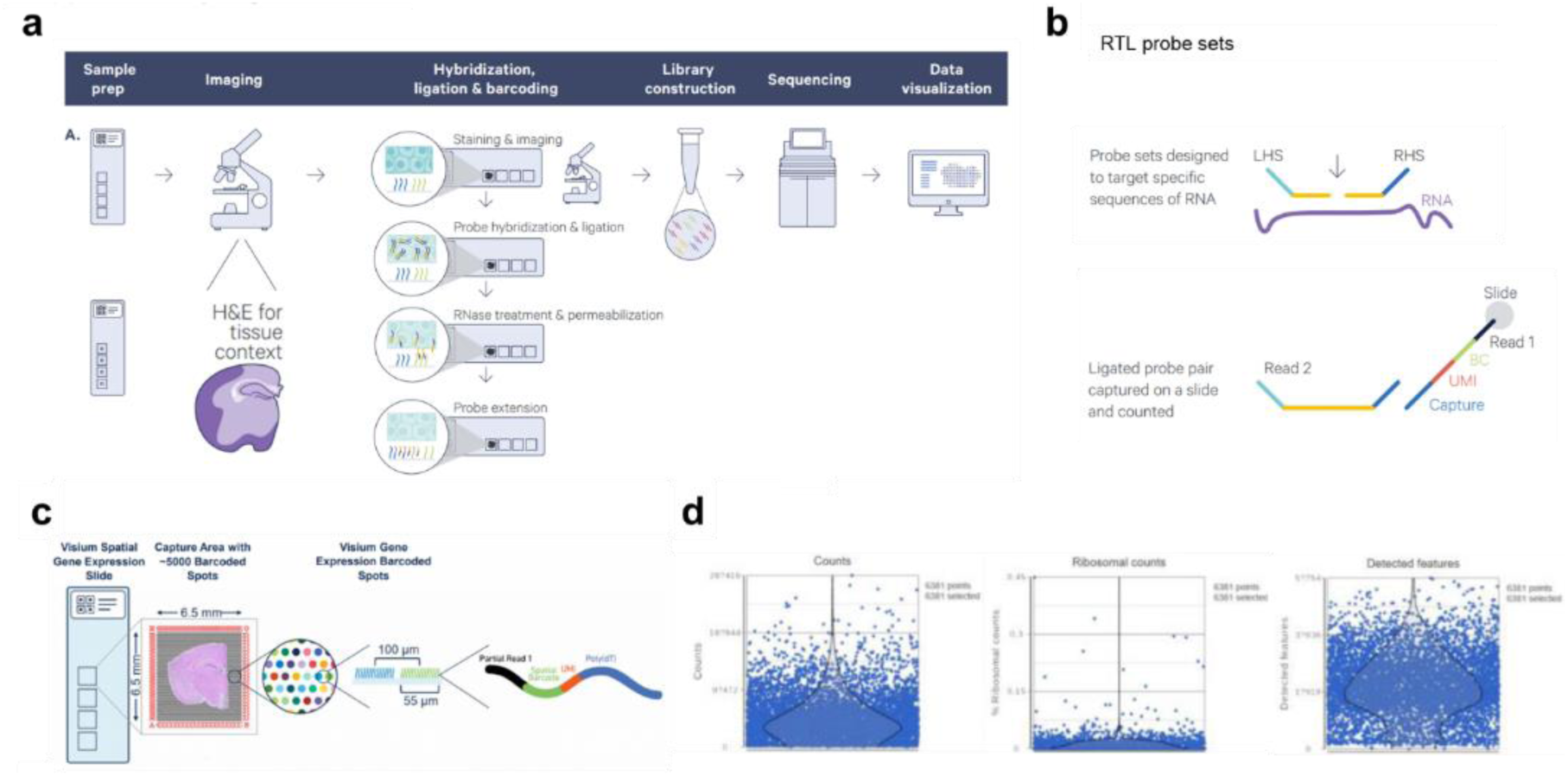
Visium technical steps and specifications. **a.** Technical steps involved in probe hybridization, extension, library construction, sequencing, and data visualization. **b.** Probe sets strategies. **c.** Visium gene expression slide capture area and spot architecture. **d.** Plots showing counts, ribosomal counts, and number of detected features per spot in Partek Flow.

**Supplementary Figure 2:**
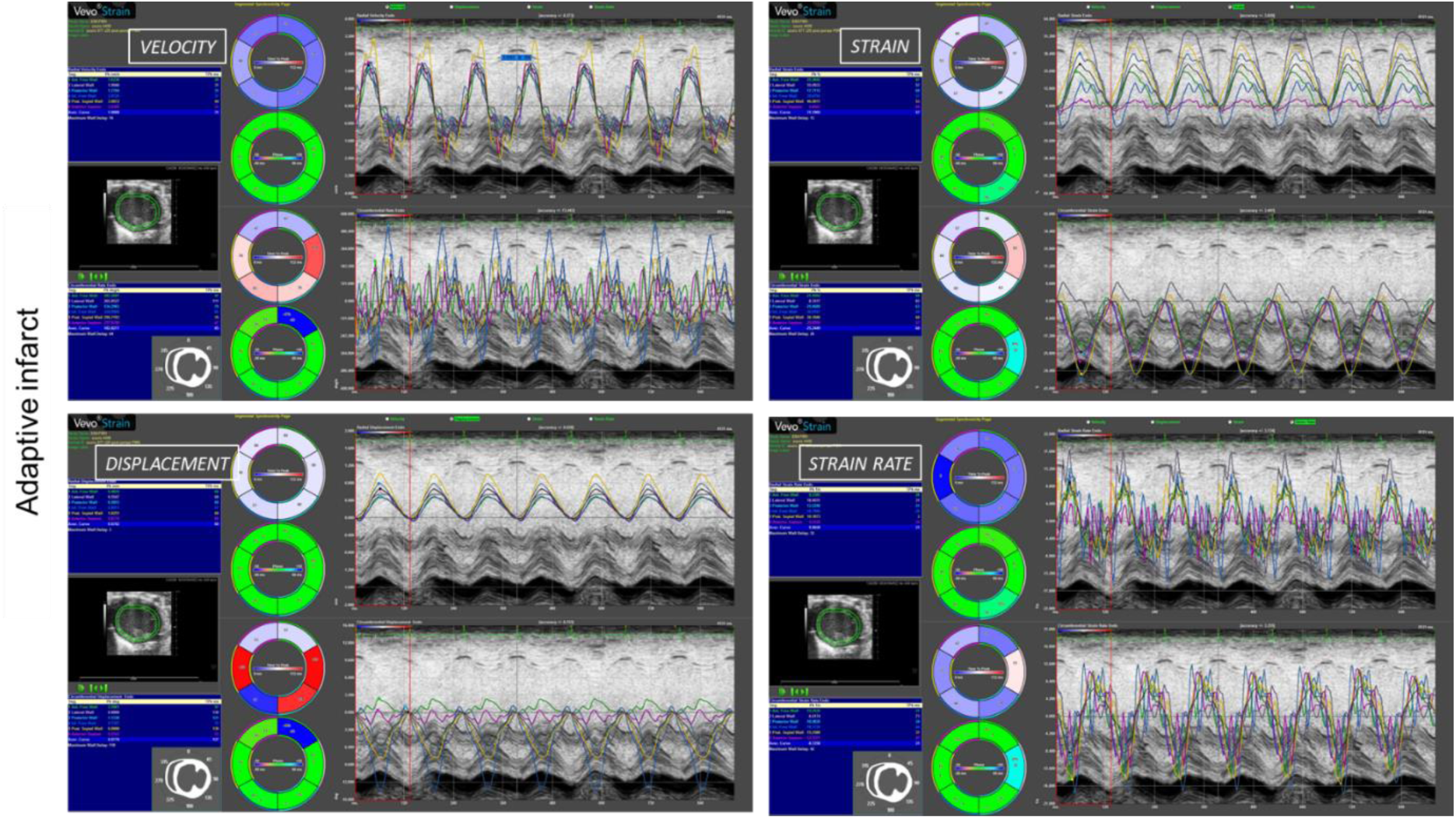
Multi parametric data showed in Vevo Strain for one adaptive infarct.

**Supplementary Figure 3:**
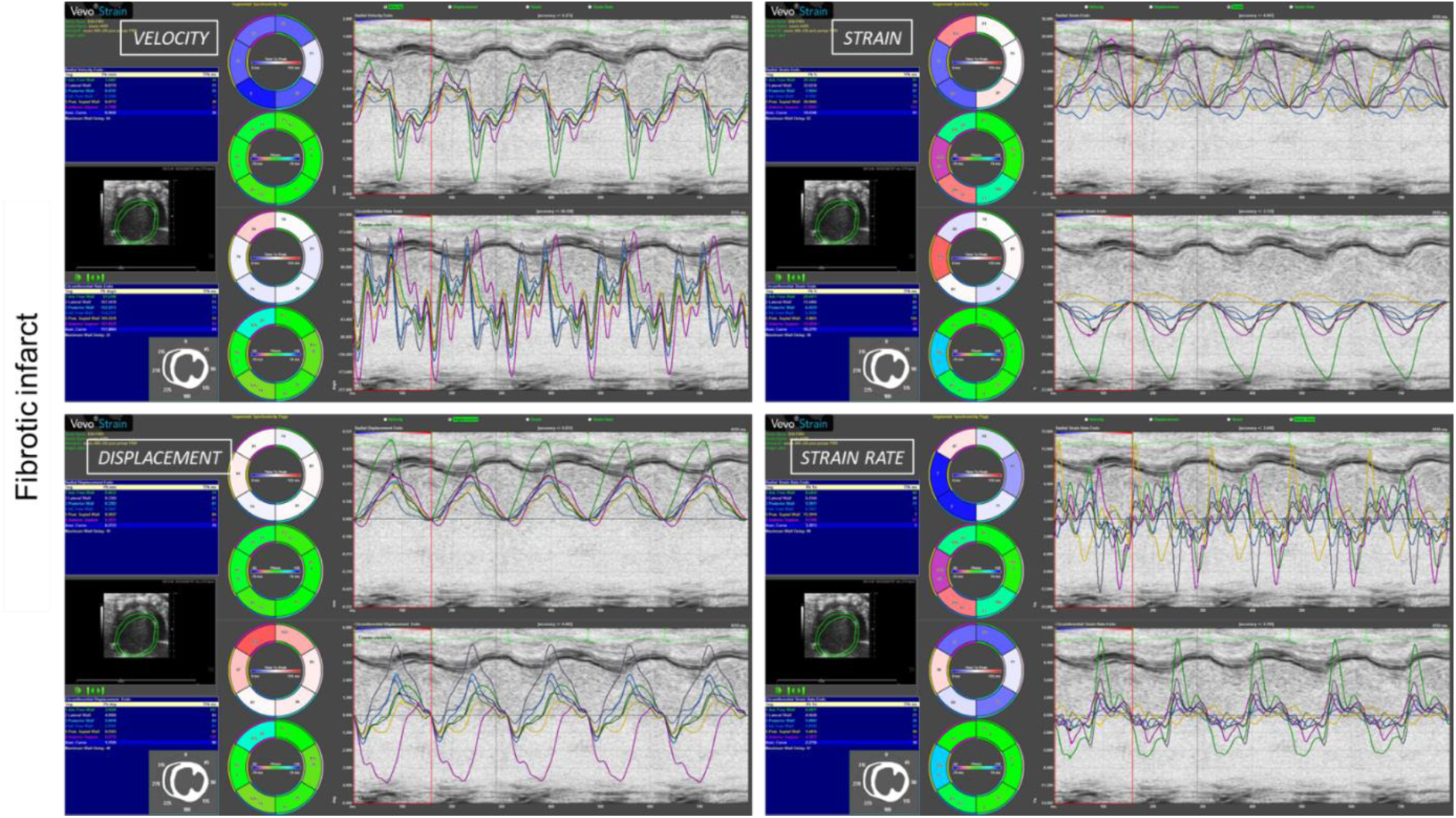
Multi parametric data showed in Vevo Strain for one fibrotic infarct.

**Supplementary Figure 4:**
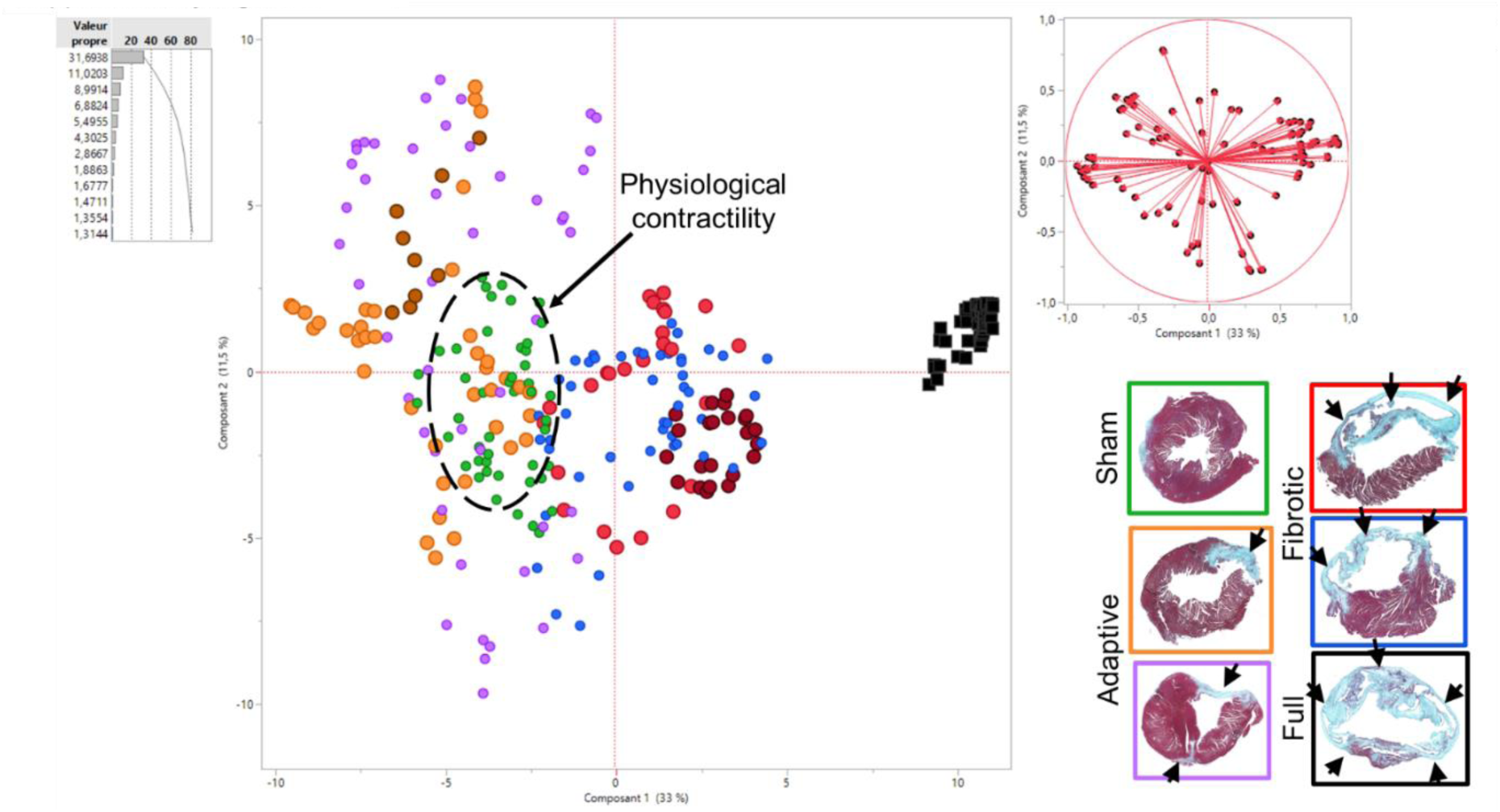
Heart motion classification using dimension reduction by principal component analysis in sham, adaptive, or fibrotic infarcts.

**Supplementary Figure 5:**
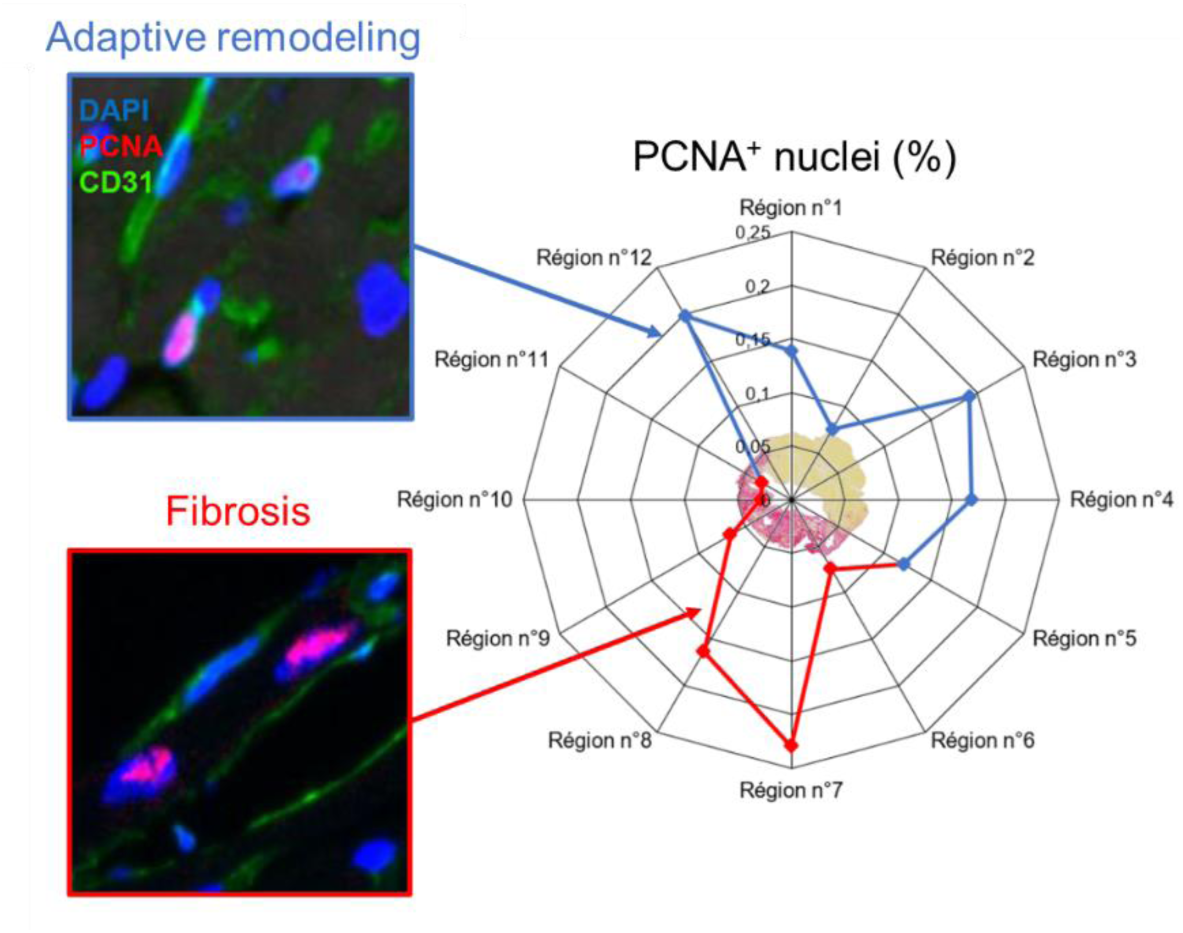
Semiquantitative evaluation of the ratio of proliferating cells (PCNA^+^) within different regions of the myocardium, in heart post-infarction (%, day 28).

**Supplementary Figure 6:**
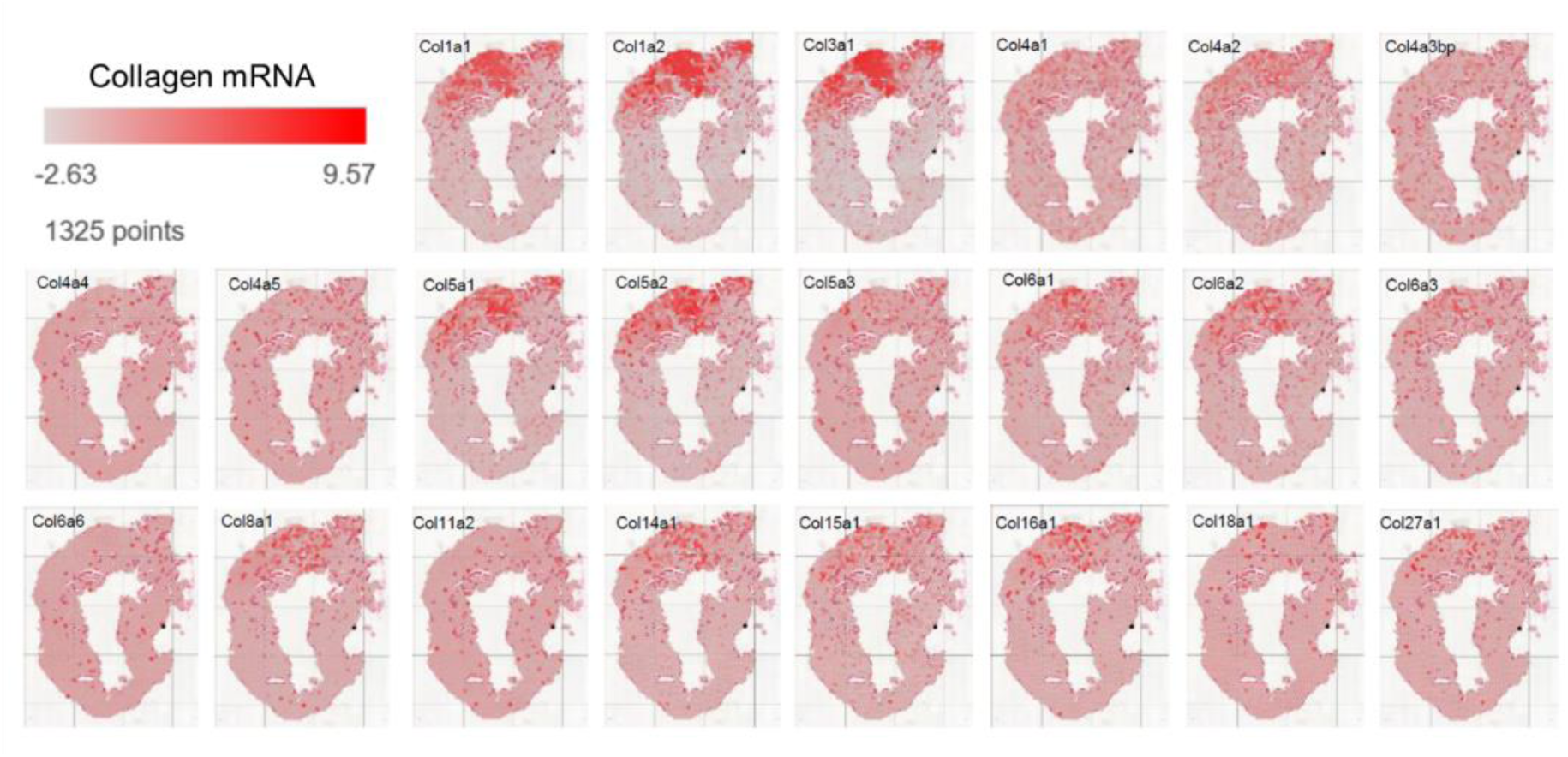
Spatial mRNA expression of various collagen family members in post-infarction heart with adaptive remodeling.

**Supplementary Figure 7:**
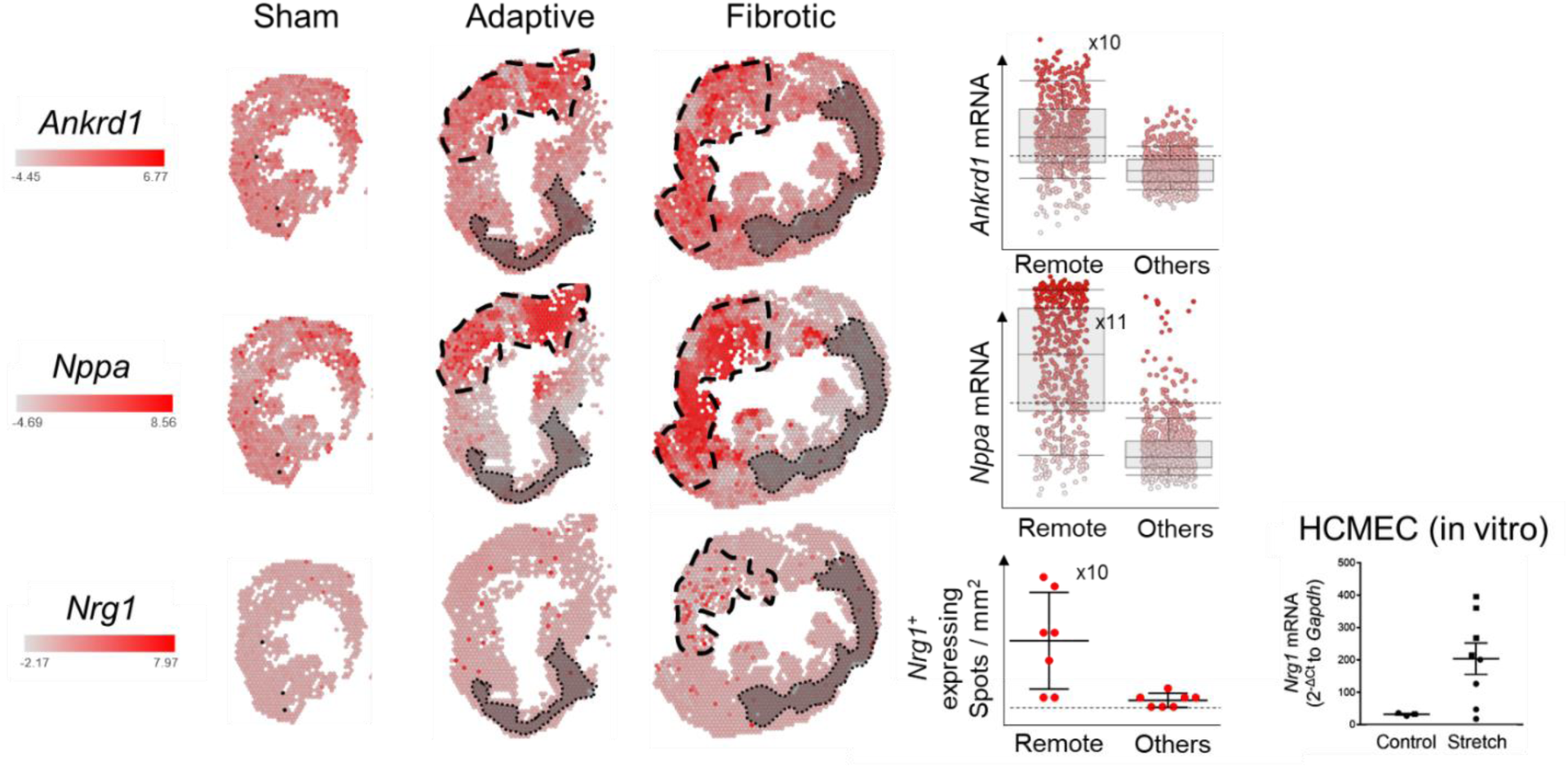
Upregulation of embryonic genes leading to hypertrophy, and *Nrg1* expression by human cardiac microvascular endothelial cells exposed to stretch (9% cyclic strain) for 8 hours.

**Supplementary Figure 8:**
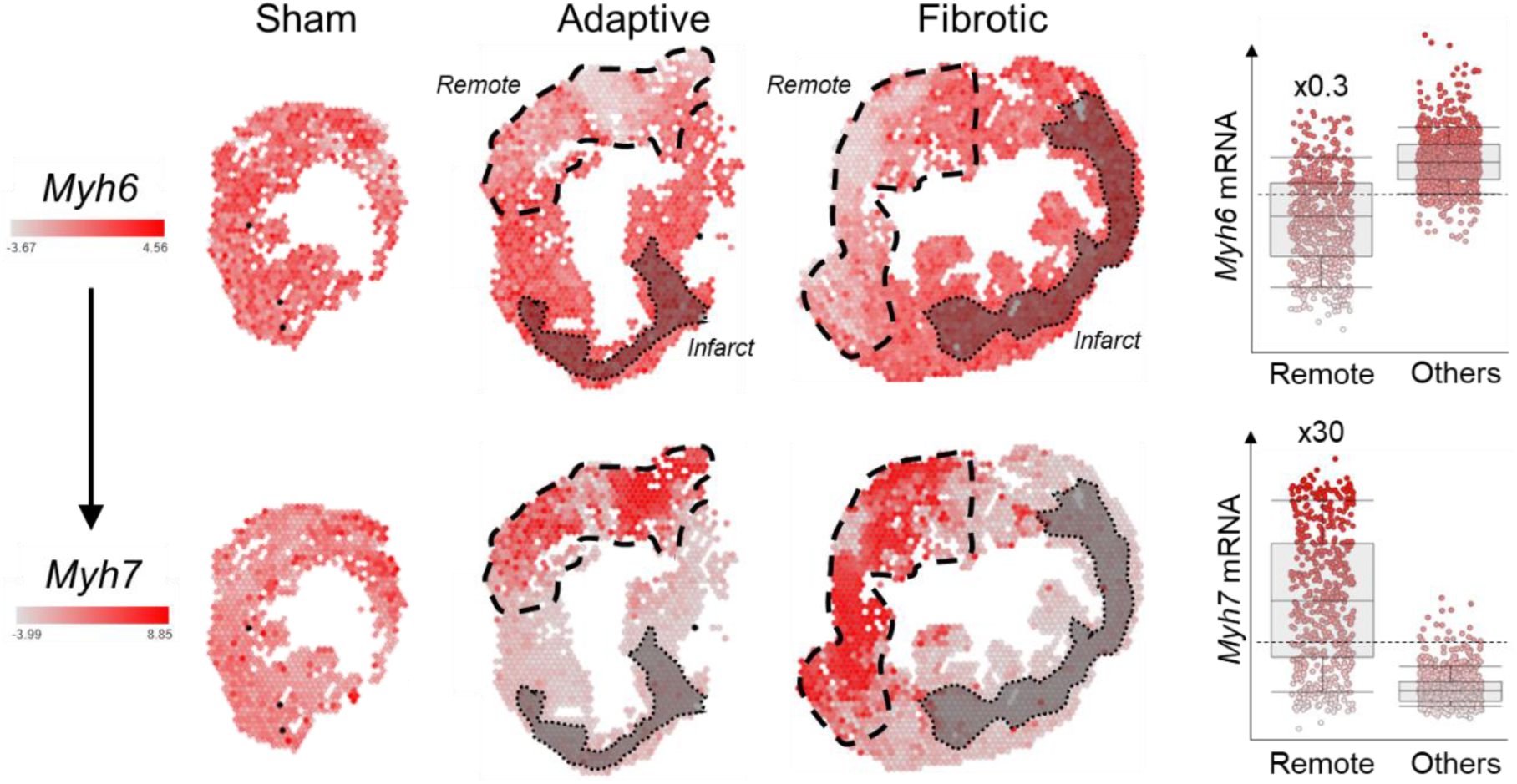
Expression inversion of the mRNA of the two major heavy chain myosin isoform *Myh6* and *Myh7*.

**Supplementary Figure 9:**
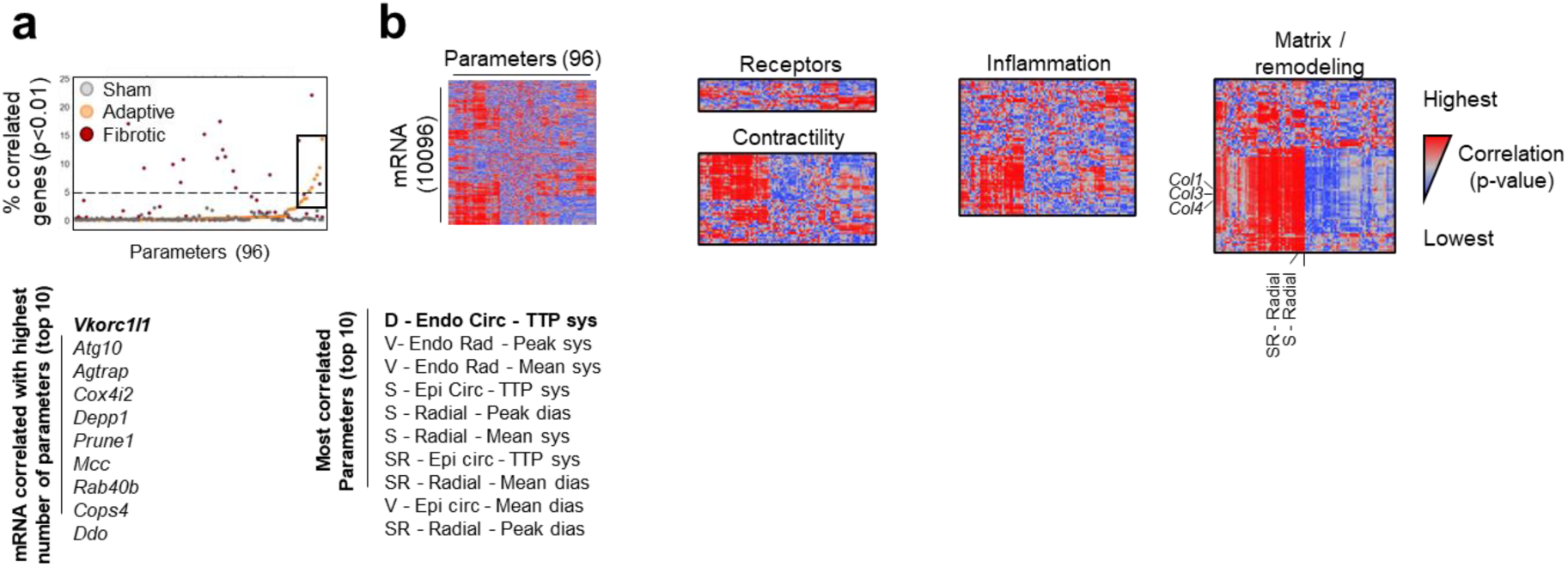
Correlation between myocardial transcriptomic analysis and mechanical forces in a post-infarction adaptive heart. Percentage of parameters significantly correlated with mRNA in an adaptive infarct. **e.** Unsupervised hierarchical clustering of the whole mRNA set according to the 96 parameters.

## References

1. Worl Health Organization, Fact sheet - Cardiovascular diseases (2017).

2. Jenca, D., et al. Heart failure after myocardial infarction: incidence and predictors. ESC Heart Fail 8, 222–237 (2021).

3. Sutton, M.G. & Sharpe, N. Left ventricular remodeling after myocardial infarction: pathophysiology and therapy. Circulation 101, 2981–2988 (2000).

4. Cottrell, C. & Kirkpatrick, J.N. Echocardiographic strain imaging and its use in the clinical setting. Expert Rev Cardiovasc Ther 8, 93–102 (2010).

5. Yuan, L.J., et al. Normal and shear strains of the left ventricle in healthy human subjects measured by two-dimensional speckle tracking echocardiography. Cardiovasc Ultrasound 12, 7 (2014).

6. Kuppe, C., et al. Spatial multi-omic map of human myocardial infarction. Nature 608, 766–777 (2022).

7. Cui, M., et al. Nrf1 promotes heart regeneration and repair by regulating proteostasis and redox balance. Nat Commun 12, 5270 (2021).

8. Muthuramu, I., Lox, M., Jacobs, F. & De Geest, B. Permanent ligation of the left anterior descending coronary artery in mice: a model of post-myocardial infarction remodelling and heart failure. J Vis Exp (2014).

9. Hafemeister, C. & Satija, R. Normalization and variance stabilization of single-cell RNA-seq data using regularized negative binomial regression. Genome Biol 20, 296 (2019).

10. Faulx, M.D., et al. Strain-dependent beta-adrenergic receptor function influences myocardial responses to isoproterenol stimulation in mice. Am J Physiol Heart Circ Physiol 289, H30–36 (2005).

11. Myagmar, B.E., et al. Adrenergic Receptors in Individual Ventricular Myocytes: The Beta-1 and Alpha-1B Are in All Cells, the Alpha-1A Is in a Subpopulation, and the Beta-2 and Beta-3 Are Mostly Absent. Circ Res 120, 1103–1115 (2017).

12. Miller, M., Koch, S.E., Veteto, A., Domeier, T. & Rubinstein, J. Role of Known Transient Receptor Potential Vanilloid Channels in Modulating Cardiac Mechanobiology. Front Physiol 12, 734113 (2021).

13. Dragun, M., Gazova, A., Kyselovic, J., Hulman, M. & Matus, M. TRP Channels Expression Profile in Human End-Stage Heart Failure. Medicina (Kaunas) 55(2019).

14. Aguettaz, E., Bois, P., Cognard, C. & Sebille, S. Stretch-activated TRPV2 channels: Role in mediating cardiopathies. Prog Biophys Mol Biol 130, 273–280 (2017).

15. Makarewich, C.A., et al. Transient receptor potential channels contribute to pathological structural and functional remodeling after myocardial infarction. Circ Res 115, 567–580 (2014).

16. Kitajima, N., et al. TRPC3 positively regulates reactive oxygen species driving maladaptive cardiac remodeling. Sci Rep 6, 37001 (2016).

17. Ferferieva, V., et al. The relative value of strain and strain rate for defining intrinsic myocardial function. Am J Physiol Heart Circ Physiol 302, H188–195 (2012).

18. Bauer, M., et al. Echocardiographic speckle-tracking based strain imaging for rapid cardiovascular phenotyping in mice. Circ Res 108, 908–916 (2011).

19. Johnson, C., Kuyt, K., Oxborough, D. & Stout, M. Practical tips and tricks in measuring strain, strain rate and twist for the left and right ventricles. Echo Res Pract 6, R87–R98 (2019).

20. Nicolosi, G.L. The strain and strain rate imaging paradox in echocardiography: overabundant literature in the last two decades but still uncertain clinical utility in an individual case. Arch Med Sci Atheroscler Dis 5, e297–e305 (2020).

